# T cell receptor (TCR) repertoire analysis reveals a highly conserved TCR repertoire in a bilateral tumor mouse model

**DOI:** 10.1101/2021.05.13.443732

**Authors:** Mikiya Tsunoda, Hiroyasu Aoki, Haruka Shimizu, Shigeyuki Shichino, Kouji Matsushima, Satoshi Ueha

## Abstract

Temporal analysis of the T cell receptor (TCR) repertoire has been used to monitor treatment-induced changes in antigen-specific T cells in patients with cancer. However, the lack of experimental models that allow temporal analysis of the TCR repertoire in the same individual in a homogeneous population limits the understanding of the causal relationship between changes in TCR repertoire and antitumor responses. A bilateral tumor model, where tumor cells were inoculated bilaterally into the backs of mice, could be used for temporal analysis of the TCR repertoire. This study examined the prerequisite for this strategy: the TCR repertoire is conserved between bilateral tumors with the same growth rate. Bilateral tumors with equivalent size and draining lymph nodes (dLNs) were collected 13 days after tumor inoculation to analyze the TCR repertoire of CD4^+^ and CD8^+^ T cells. The tumor-infiltrating T cell clones were highly conserved between the bilateral tumors, and the extent of clonal expansion was equivalent. In addition, the similarity between the bilateral tumors was equivalent to heterogeneity on one side of the tumor. The similarity of the TCR repertoire in the bilateral dLNs was markedly lower than that in the tumor, suggesting that tumor-reactive T cell clones induced independently in each dLN integrated during recirculation and then infiltrated the tumor. These findings suggest that our bilateral tumor model is suitable for temporal monitoring of the TCR repertoire to evaluate temporal and treatment-induced changes in tumor-reactive T cell clones.

**Significance Statement:** The bilateral subcutaneous tumor model, where tumor cells were inoculated bilaterally into the backs of mice, is a promising model for temporal analysis of the antitumor response in cancer immunotherapy. This study demonstrated a highly conserved TCR repertoire in bilateral tumors and provided the basis for using a bilateral tumor model for evaluating temporal and treatment-induced changes in tumor-responsive T cell clones in individual mice. In humans, accurate statistical analysis is hampered by various background factors, including cancer type and stage and history of treatment. Moreover, temporal tumor biopsy in patients is highly invasive. Our bilateral tumor model overcomes these clinical issues and is expected to be a valuable tool for the development of novel immune monitoring and therapeutic strategies.

## Introduction

Immune checkpoint inhibitors (ICIs) have a significant therapeutic effect in some cancers and have become an important pillar of cancer treatment in recent years (1, 2). However, the response rate to ICI monotherapy is less than 30% for most types of cancer, and ICIs occasionally cause severe immune-related adverse effects in some patients (3, 4). Thus, the development of reliable biomarkers that represent tumor-specific immune responses and stratify the responder and non-responder at an early stage is essential to optimize the usage of ICIs (5, 6).

Because ICIs suppress tumor growth by enhancing the proliferation and activation of tumor-specific T cells (7), the efficacy of ICIs is closely associated with the strength of tumor-specific T cell responses. These tumor-specific T cells are composed of various tumor-reactive T cell clones with different specificities to tumor-associated antigens (8, 9). Antigen specificity of T cell clones is determined by their T cell receptors (TCRs), which are generated by V(D)J recombination in the thymus and are incredibly diverse (10, 11). Therefore, global analysis of the collection of TCRs using next-generation sequencing, that is TCR repertoire analysis, can now be applied to monitor tumor-specific T cell responses in patients receiving ICIs.

Several studies have reported the features of the TCR repertoire in mice treated using immunotherapy. Philip and Rudqvist reported enhanced clonal expansion of tumor-infiltrating T cells in mice treated with an anti-cytotoxic T lymphocyte-associated protein 4 monoclonal antibody (12, 13). We reported that CD8^+^ T cell clones with a TCR identical to tumor-infiltrating T cells could be detected in the peripheral blood and draining lymph node (dLN) of tumor-bearing mice (14). Based on the cancer-immunity cycle, where tumor-specific T cells are primed in the dLN and infiltrate into the tumor via lymph–blood circulation (15), these “overlapping” T cell clones seem to reflect the mobilization of T cells into antitumor responses. Consistent with this hypothesis, the diversity and total frequency of CD8^+^ T cell clones that overlapped between the blood and tumor increased after depletion of CD4^+^ cells, the degree of which was correlated with antitumor effects in mice and humans (14, 16). To validate whether these features of the TCR repertoire are predictive of the antitumor effect of ICIs, an appropriate analysis of the temporal or treatment-induced changes in TCR repertoire is needed. However, temporal analysis of the TCR repertoire of tumor-reactive T cells has not been reported in murine tumor model, owing to the difficulty in sampling from an individual mouse.

Recently, Zemek and Chen reported that the immune microenvironment is similar in bilateral tumor models with comparable tumor growth and that it can be applied to temporal analysis of antitumor immune responses (17, 18). To apply this method for temporal analysis of the TCR repertoire, there are important prerequisites: (1) the TCR repertoires of the left and right tumors show similar characteristics; and (2) the same tumor-reactive T cell clones infiltrate into the bilateral tumors. In this study, we investigated whether the bilateral tumor model is suitable for examining the temporal responses of tumor-reactive T cells in individual mice.

## Results

### T cell repertoire of bilateral tumor exhibits similar characteristics

We used the bilateral tumor model to establish an experimental system for evaluating temporal and treatment-induced changes in tumor-reactive T cell clones. To achieve this goal, the growth rate and clonal T cell responses in bilateral tumors must be similar in individual mice. Therefore, we first examined whether the Lewis lung carcinoma (LLC) tumors inoculated bilaterally into the back of individual mice grew symmetrically. Intratumoral injection of Evans Blue verified that the brachial lymph node became a dLN in the subcutaneous tumors (Fig. 1A). The growth curves of bilateral tumors were symmetrical in individual mice, suggesting that an equivalent antitumor response occurred on both sides of the tumor (Fig. 1B).

**Figure. 1.**
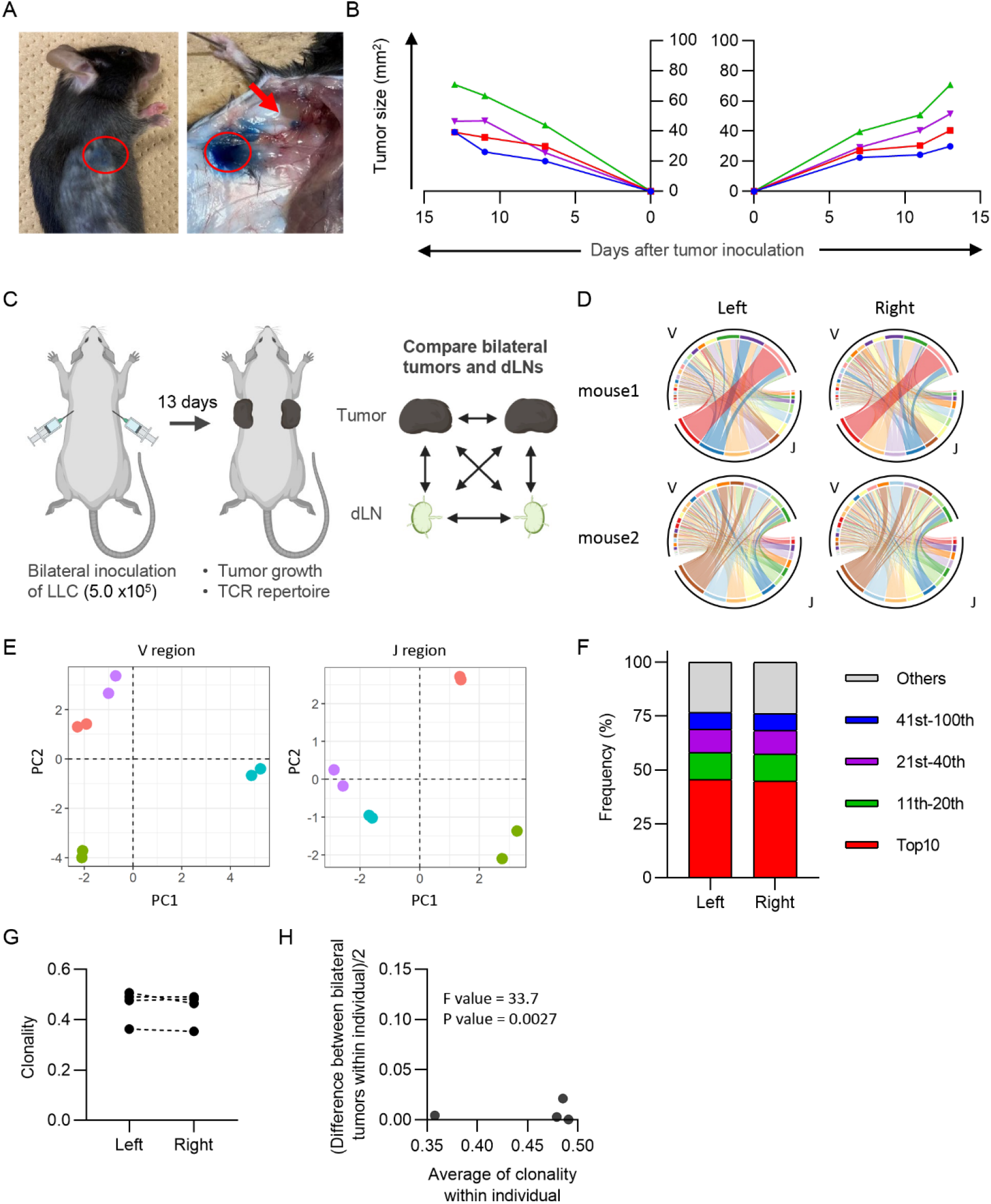
Characteristics of CD8^+^ T cell repertoire in the bilateral tumor. (A) Visualization of draining lymph node (dLN) in our bilateral tumor model. 1% Evans Blue dye was injected into the tumor (red circle) 30 min prior to euthanasia. Brachial LN (arrow) was stained by Evans Blue, indicating this LN became a dLN. (B) Growth curve of the tumor inoculated bilaterally on the backs. Lines of the same color indicate left and right tumors of the same mouse (n = 4). (C) Experimental design. (D) V/J segment usage plots of the bilateral CD8^+^ T cell repertoires. Ribbons connecting the V and J segments are scaled by the corresponding V/J pair frequency. (E) Principal component analysis of V and J segment usage of tumor-infiltrating CD8^+^ T cells. Points of the same color indicate left and right tumors of the same individual. (F) Frequency of abundant CD8^+^ T cell clones in the tumor. CD8^+^ T cell clones in the left and right tumor were categorized into five classes based on their rank in each repertoire: top 10, 11^th^-20^th^, 21^st^-40^th^, 41^st^-100^th^, and others. The total frequency of clones in each class is shown (n = 4). (G) Clonality of the CD8^+^ T cell repertoire of the left and right tumor. (H) Homoscedasticity plot for variance of clonality in individual mice. The X-axis represents the average clonality of bilateral tumors in each individual. The Y-axis represents variance of clonality between different bilateral tumors within individuals. Sum of squares within mouse = 9.7 × 10^-4^; sum of squares between mice = 2.4 × 10^-2^; degree of freedom within mouse = 4; degree of freedom between mice = 3.

Next, we investigated the equivalency of T cell clonal responses inside the bilateral tumors. CD4^+^ and CD8^+^ T cells from the tumor and CD4^+^ CD44^hi^ and CD8^+^ CD44^hi^ T cells from dLN were sorted 13 days after tumor inoculation, and their TCR repertoires were analyzed (Fig. 1C and SI Appendix Fig. S1A). Variable-Joining (V/J) segment usage of TCRβ, which is commonly used to characterize individual repertoire, was highly similar between bilateral tumors in the same mouse, but variable among mice (Fig. 1D and E). The proportion of the most abundant T cell clones was also similar between bilateral tumors, suggesting that they contained an equivalent number of expanded clones (Fig. 1F). Consistently, the clonality of the CD8^+^ T cell repertoire, which represents the extent of clonal expansion, was equivalent between the bilateral tumors (Fig. 1G). In addition, the difference in clonality between the bilateral tumors of the same mouse was significantly smaller than that between different mice (Fig. 1H). A similar trend was observed for the CD4^+^ T cell repertoire (SI Appendix Fig. S1). These results demonstrated that the global features of the TCR repertoire were similar between the bilateral tumors.

### The majority of the T cell repertoire was composed of shared clones with a similar extent of expansion in bilateral tumors

Highly similar features of the TCR repertoire between bilateral tumors suggested that T cell clones present on one side also exist on the other side. To examine this, we identified T cell clones that overlapped between the bilateral tumors or different individuals (Fig. 2A). The clones shared between bilateral tumors accounted for approximately 80% of the CD8^+^ T cell repertoire. However, T cell clones overlapping between different mice covered only approximately 3%. This result indicated that most of the tumor-infiltrating CD8^+^ T cell clones were commonly present on both sides of the tumor.

**Figure. 2.**
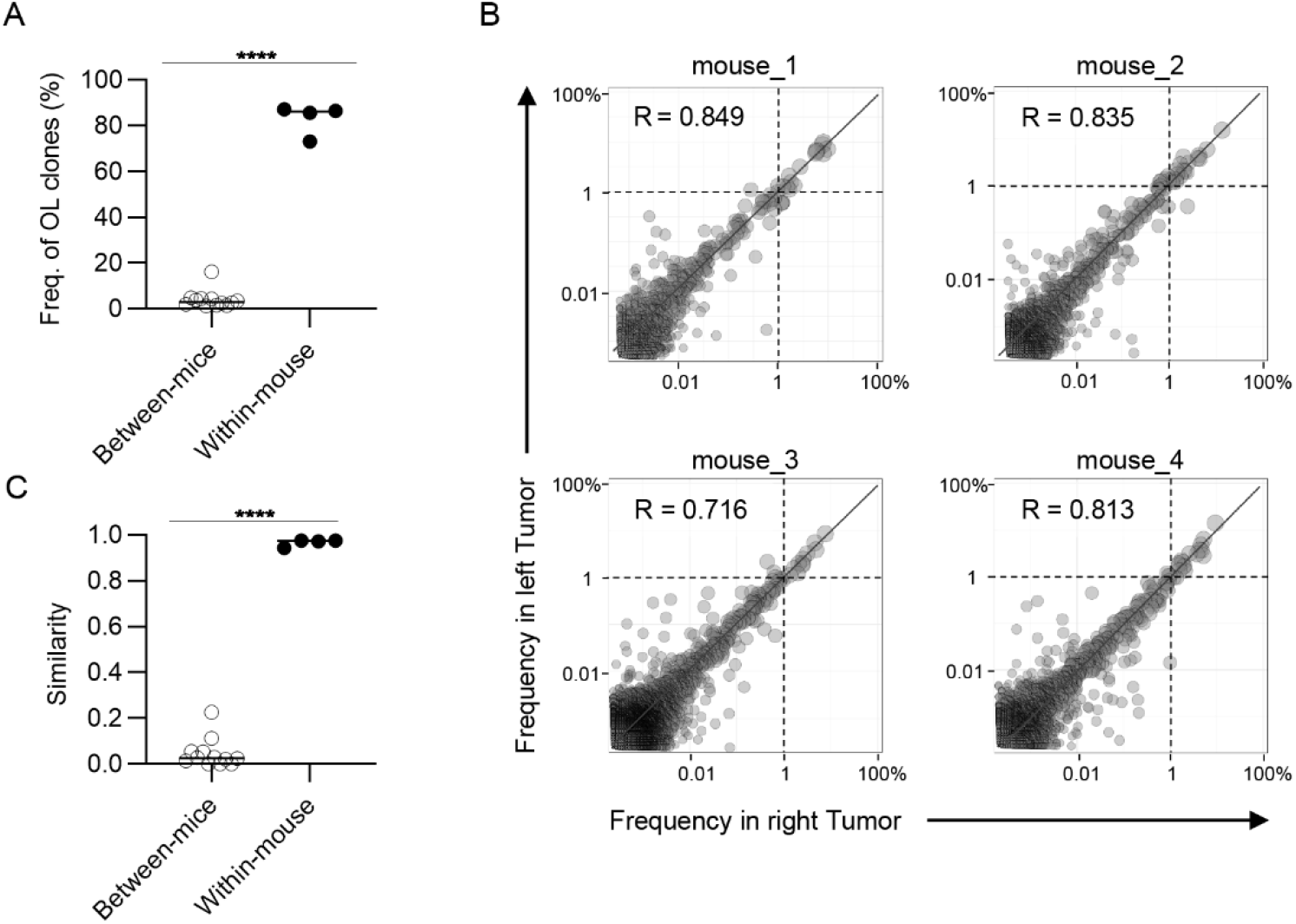
Clonal composition of CD8^+^ T cell repertoires in bilateral tumors. (A) Comparison of the frequency of overlapping clones in tumors between and within mice (between mice, n = 4 × 3; within mice, n = 4). (B) Scatter plot of CD8^+^ T cell clones from bilateral tumors. Each plot represents a single clone with indicated frequency in the left (X-axis) and right tumors (Y-axis). The dotted line indicates a frequency of 1%. (C) Comparison of the similarity of the tumor repertoires between and within mice. Mean; Two-sided unpaired Student’s t-test (A and C); **** P ≤ 0.0001.

Next, we examined whether these overlapping clones expanded equally in bilateral tumors. To this end, we depicted the frequency of each overlapping clone within the right (x-axis) and left (y-axis) tumors as a scatter plot (Fig. 2B). The frequency of clones that had expanded to more than 1% was almost equal between the left and right sides. There were some clones whose frequency differed between the left and right sides, but the frequency of these clones was relatively low. Consistent with this finding, the Morisita-Horn similarity index, which reflects the similarity of two T cell repertoires considering the frequency of each shared clone (19), was over 0.9 between the bilateral tumors, whereas it ranged from 0.05 to 0.3 among the tumors in different individuals (Fig. 2C). The CD4^+^ T cell repertoire also showed similar tendencies (SI Appendix Fig. S2). These results suggest that the majority of CD8^+^ T cell clones in tumors respond to tumor antigens shared between the bilateral tumors. Altogether, these data showed that the majority of the T cell repertoire of bilateral tumors was composed of shared clones, and these clones expanded to a similar extent in the left and right tumors. Thus, the clonal T cell responses on one side may reflect those on the other side in our bilateral tumor model.

### Similarity of TCR repertoires between bilateral tumors was equivalent to the similarity within the tumor

Previous studies have shown that there is intratumoral heterogeneity of the TCR repertoire (20). When extrapolating our bilateral tumor model to a clinical situation, where antitumor immune responses are longitudinally monitored by tumor biopsy, it is important to determine whether the difference in TCR repertoire between the bilateral tumors could be considered equivalent to that within one side of the tumor. To examine this, we divided each tumor into two pieces at the time of collection and analyzed the TCR repertoire independently to estimate intra-tumoral TCR heterogeneity. We then compared the similarity of the TCR repertoire within one side of the tumor and between bilateral tumors (Fig. 3A). The total frequency of CD8^+^ overlapping clones between the left and right tumors was approximately 80%, which was equivalent to the frequency of overlapping clones within one side (Fig. 3B). Additionally, scatter plots depicting the frequency of each overlapping clone showed that the variance in frequency of overlapping clones between bilateral tumors was equivalent to that within the tumor (Fig. 3C). Consistently, there was no significant difference in the similarity of the TCR repertoire between the tumor and within the tumor (Fig. 3D). A similar tendency was observed for the CD4^+^ T cell repertoire (SI Appendix Fig. S3). These data demonstrated that the differences in TCR repertoires between the bilateral tumors were equivalent to the TCR heterogeneity within each tumor. This also suggested that our bilateral tumor experiments could be considered as an experimental model for temporal monitoring of TCR repertoire using sequential tumor biopsy.

**Figure. 3.**
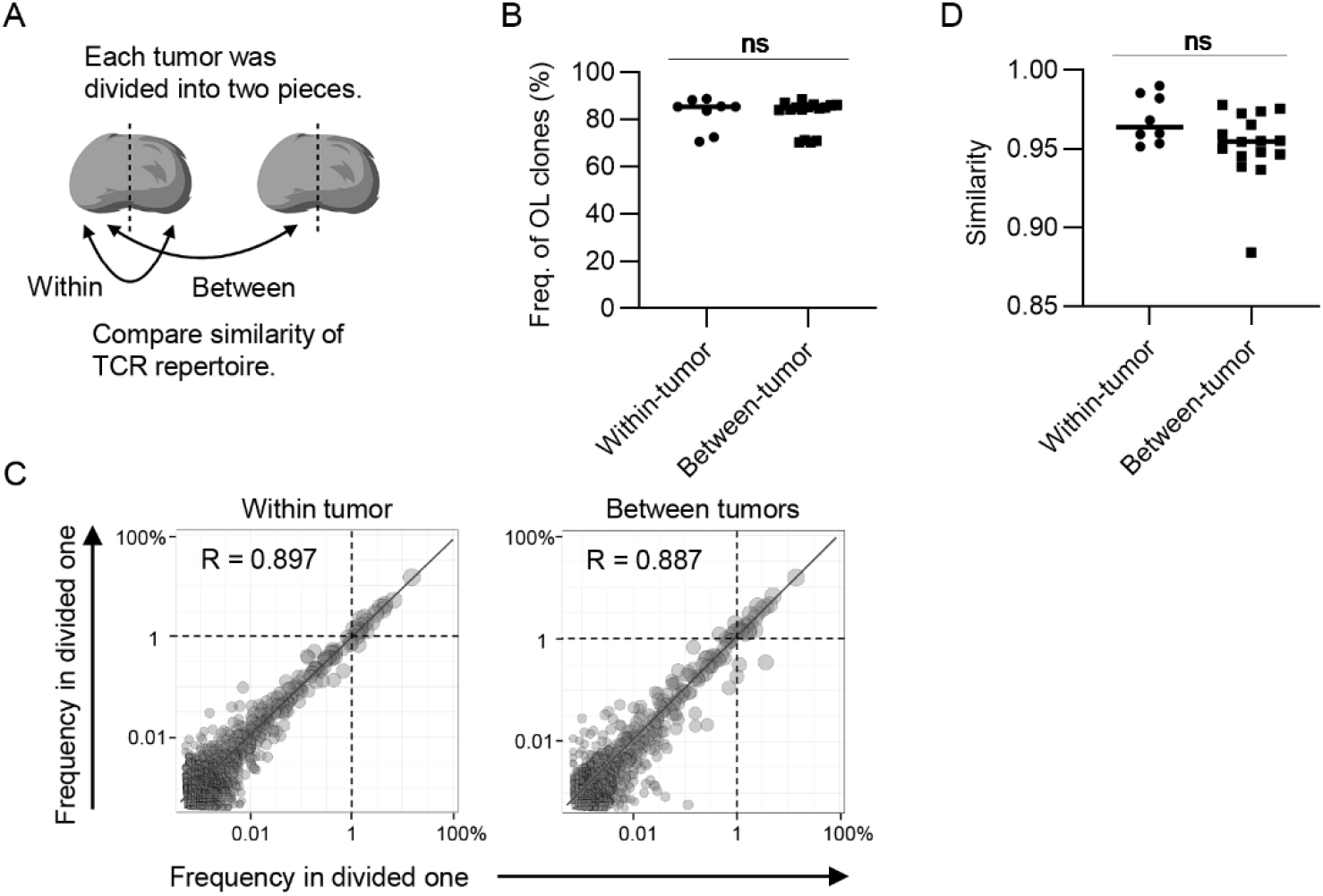
The extent of difference of CD8^+^ T cell repertoires between bilateral tumors. (A) Schematic diagram of the analysis. (B) Comparison of the frequency of the clones overlapped between divided same side tumors and bilateral tumor fragments (n = 4). (C) Scatter plot of CD8^+^ T cell clones from bilateral tumors. Each plot represents a single clone with indicated frequency in each tumor fragment. The dotted line indicates a frequency of 1%. (D) Comparison of the similarity of the tumor repertoires between divided same tumors and bilateral tumor fragments. (B and D) Within-tumor, n = 4 × 2; between-tumor, n = 4 × 4; Mean; Two-sided unpaired Student’s t-test; ns, non-significant.

### Proportional infiltration of T cell clones into bilateral tumors contributed to the similarity of TCR repertoires

Tumor-reactive clones that expand in the bilateral dLNs exit from the efferent lymph, enter the blood circulation via the thoracic duct, and eventually form a blood repertoire. Thus, it is hypothesized that the tumor-reactive clones induced on one side of the dLNs infiltrate into the bilateral tumors at equal probability through the circulation and then proliferate *in situ* at the same rate (Fig. 4A). To explore this idea, we investigated whether the frequency of overlapping clones between the tumor and its contralateral dLN was equivalent to that of overlapping clones between the ipsilateral ones (Fig. 4A). The total frequency of overlapping clones between contralateral dLNs in the tumor was almost the same as that between the ipsilateral ones, and this result supported our idea (Fig. 4B).

**Figure. 4.**
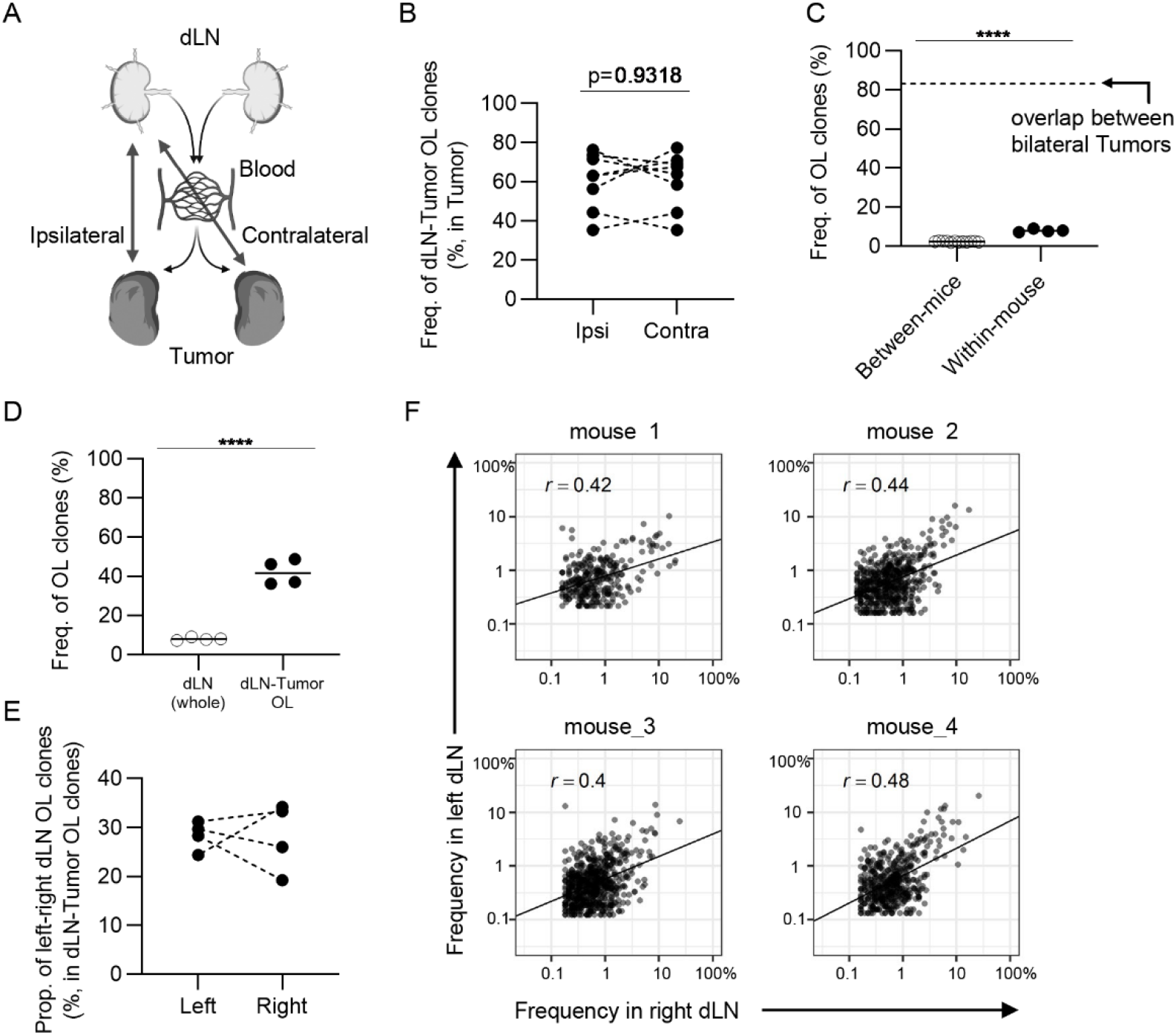
The similarity of CD8^+^ T cell repertoires in the bilateral dLNs. (A) Schematic diagram of the hypothesis that T cell clones induced in dLN infiltrate into tumors evenly through blood circulation. (B) The frequency of the clones overlapped between the dLN and its ipsilateral and contralateral tumor. Ipsi, ipsilateral; Contra, contralateral. n = 4 × 2. (C) Comparison of the frequency of overlapping clones in dLN between and within mice (between mice, n = 4 × 3; within mice, n = 4). Dotted line indicates the frequency of clones overlapped between bilateral tumors within the mouse. (D) Comparison of the frequency of left–right overlapping clones within whole dLNs and dLN-tumor overlapping repertoire (n = 4). (E) The proportion of left–right overlapping clones within the dLN-tumor OL repertoires. (F) Scatter plot of left–right overlapping clones within the dLN-tumor OL repertoires. Each plot represents a single clone with indicated frequency in each dLNs. Mean; Two-sided unpaired Student’s t-test; ns, non-significant. **** P ≤ 0.0001.

However, it was unclear whether similar clones were induced in bilateral dLNs. To address this query, we examined the frequency of overlapping clones in the bilateral dLNs and found that it was lower than 10%, which was substantially lower than the overlap between bilateral tumors, although it was higher than the overlap between individuals (Fig. 4C). Because the dLN CD8^+^ CD44^hl^ T cells may contain a substantial proportion of clones not associated with the tumor, we next examined the TCR repertoire of bilateral dLNs that overlapped with either of the bilateral tumors to enrich tumor-associated clones. In dLN-tumor overlapping clones in the bilateral dLNs, the frequency of left–right overlapping clones between bilateral dLNs increased to approximately 40 % (Fig. 4D), and the proportion of left–right overlapping clones was approximately 20-40% (Fig. 4E). In other words, more than 50% of dLN-tumor overlapping clones were induced only in one of the dLNs. Finally, we investigated whether these dLN-tumor overlapping clones were equally expanded in the bilateral dLNs. Using scatter plots, we showed that the correlation in frequency in dLN-tumor overlapping clones was moderate between the bilateral dLNs (0.4 < r < 0.48; Fig. 4F), compared to the correlation between the bilateral tumors (0.71 < r < 0.85; Fig. 2B). The frequency of CD4^+^ T cell clones overlapping between bilateral dLNs was higher than that of CD8^+^ T cell clones, and the CD4^+^ T cell repertoire also showed similar tendencies (SI Appendix Fig. S4).

Collectively, these results suggested that the tumor-reactive T cell clones induced in the bilateral dLNs were only moderately conserved, and the degree of expansion of the shared clones differed between the bilateral dLNs. Therefore, the proportional infiltration of T cell clones into bilateral tumors through blood circulation contributed to the conserved TCR repertoire.

## Discussion

The bilateral subcutaneous tumor model, where tumor cells were inoculated bilaterally into the backs of mice, is a promising model for temporal analysis of the antitumor response in cancer immunotherapy. In this study we examined the prerequisite for this strategy: the TCR repertoire is conserved between bilateral tumors with the same growth rate. We found that bilateral tumors with similar growth rates contained a highly conserved CD8^+^ and CD4^+^ T cell repertoire in our experimental model. Interestingly, the TCR repertoires in bilateral dLNs were less conserved than those in bilateral tumors. These results suggest that T cell clones induced in the bilateral dLNs eventually integrated into a blood repertoire, proportionally infiltrated the bilateral tumors, and proliferated in situ at a similar rate. These findings provide the basis for analyzing temporal and treatment-induced changes in tumor-reactive T cell clones using a bilateral tumor model in mice.

Previous reports have examined the relationship between the immune microenvironment before ICI treatment and the antitumor response in bilateral tumor models; Zemeck et al. reported that in bilateral tumor models, the transcriptional signature of tumors before ICI treatment was different between responders and non-responders (17). Chen and colleagues analyzed tumor-infiltrating CD8^+^ T cells in bilateral tumor models treated with ICI and reported that non-responders had an exhaustion signature and responders had an activation signature (18). All of these reports suggest that similar immune and antitumor responses are induced in bilateral tumors. Our finding of a highly conserved TCR repertoire in bilateral tumors may explain why similar immune responses and gene expression are induced bilaterally.

Although we did not test other mouse tumor models, our bilateral tumor model would be applicable in other tumor models, because the conserved TCR repertoire between the bilateral tumors seems to be dependent on anatomical factors that are absolutely conserved among individuals. Notably, the difference in tumor size may alter the tumor microenvironment, such as the concentrations of chemo-attractants and vascularity, between bilateral tumors, and it may decrease the similarity in TCR repertoire between the bilateral tumors. Verifying the bilateral symmetry of tumor growth or therapeutic response is necessary to evaluate the temporal changes in the tumor-reactive T cell repertoire using this model.

In humans, patient background data, such as cancer type and stage and history of treatment are associated with prognosis (21). Thus, it was difficult to obtain a large cohort of homogeneous patients for statistical analysis. Moreover, temporal tumor biopsy in patients is highly invasive. Considering these difficulties, clinical studies to validate the relationship between TCR repertoire and antitumor effects are limited. We believe that our bilateral tumor mouse model will overcome these barriers and provide valuable suggestions for clinical research.

A possible application of this bilateral tumor model is to investigate the features of the TCR repertoire that can predict the therapeutic response to ICIs. Considering that the antitumor responses following ICI therapy vary even among mice with the same genetic background, the specific features of the TCR repertoire may reflect the variance of antitumor immune responses among syngeneic mice. We plan to examine the hypothesis that a large amount of dLN-tumor repertoire overlap before treatment would predict a better therapeutic response to ICIs using the bilateral tumor model. Clinically, we observed that an increased frequency of tumor-blood overlapping clones in blood CD8+ T cells before treatment was associated with a favorable clinical response to PD-1 blockade in gastrointestinal cancer (22). In addition, the bilateral tumor model enables temporal tracking of endogenous T cell clones in the same mouse, while tumor progression or therapeutic effects continue. We expect that temporal analysis of T cell clones within bilateral tumors will reveal the kinetics of expansion and contraction of each T cell clone and the process of exhaustion in the tumor.

Overall, this study reported the TCR repertoire analysis of bilateral tumor models, which enables the evaluation of temporal and treatment-induced changes in the tumor-reactive T cell clones. We believe that this novel experimental system will deepen our understanding of the clonal responses of tumor-reactive T cells.

## Materials and Methods

### Mice and cell line

Eight-week-old female C57BL/6 mice were purchased from Sankyo Labo service corporation inc. (Tokyo, Japan). Lewis lung carcinoma (LLC) was originally provide from the Nihonkayaku (Tokyo, Japan).

### Tumor inoculation

LLC cells (5 × 10^5^ cells /mouse) were inoculated subcutaneously (s.c.) into the bilateral backs of C57BL/6 mice. Tumor diameter was measured twice a week and used to calculate tumor volume (mm) according to the following formula [(major axis; mm) x (minor axis; mm)]. In some mice, 1% Evans Blue dye was injected into the tumor 30 min prior to sacrifice to determine the dLN in our model. All animal experiments were conducted in accordance with institutional guidelines with the approval of the Animal Care and Use Committee of the Tokyo University of Science.

### Flow cytometry and Cell Sorting

Intravascular leukocytes were stained by intravenous injection of FITC-conjugated mAb (3 μg/mouse) against CD45.2 (clone 104) three minutes before sacrifice (23). Tumor was equally divided into two parts and processed individually. Each tumor was cut into small fragments and digested for 45 minutes at 37°C with 0.1% collagenase (FUJIFILM Wako, Osaka, Japan). The cells were then subjected to density separation with 40% Percoll PLUS (Cytiva, Marlborough, MA) and leukocytes were recovered from the bottom layer. ACK buffer was used to lyse red blood cells. The extracted dLN was cut into small fragments and mashed on a cell strainer. The cell number was determined using Flow-Count fluorospheres (Beckman Coulter, San Diego, CA) and a CytoFLEX flow cytometer (Beckman Coulter). Cells were then stained with a mix of Fc Block (anti-mouse CD16/CD32 mAb; clone 2.4G2, BioXcell) and fluorophore-conjugated anti-mouse mAbs as indicated in Table S1. After enrichment of T cells with magnetic separation, CD8^+^ and CD4^+^ T cells from the tumor and CD8^+^ CD44^hi^ and CD4^+^ CD44^hi^ T cells from the dLN were sorted using FACS Aria II or Aria III (BD Biosciences, San Jose, CA). Nonviable cells were excluded from the sorting gate based on forward and side scatter profiles and propidium iodide staining. intravascular staining-CD45.2 positive cells were also excluded. Purity of sorted cells was always over 95%. Data were analyzed using FlowJo software (version 10.5.3; BD Biosciences).

### TCR library construction and sequencing

TCR libraries were prepared on purified T cells lysed in lysis buffer [1% LiDS, 100 mM Tris-HCl (pH 7.5), 500 mM LiCl, and 10 mM EDTA]. PolyA RNAs were isolated according to a previous report with some modifications (GSE110711).To perform reverse transcription and template-swiching, beads were then suspended in 10 μL of RT mix [1× First Strand buffer (Thermo Fisher Scientific), 1 mM dNTP, 2.5 mM DTT (Thermo Fisher Scientific), 1 M betaine (Sigma-Aldrich), 9 mM MgCl2, 1 U/μL RNaseIn Plus Rnase Inhibitor (Promega, Madison, WI), 10 U/μL Superscript II (Thermo Fisher Scientific), and 1μM of i5-TSO], and incubated for 60 min at 42°C and immediately cooled on ice. Beads were washed once with B&W-T buffer [5 mM Tris-HCl (pH 7.5), 1 M NaCl, 0.5 mM EDTA, and 0.1% Tween-20], and once with Tris-HCl (pH 8.0). To amplify the TCR cDNA containing complementarity determining region 3 (CDR3), nested PCR of the TCR locus was performed as follows. Beads containing cDNA was resuspended with the 25 μL of first PCR mixture [0.4 μM of primers (i5, Trac_ex, and Trbc_ex), and 1x KAPA Hifi Hotstart ReadyMix (KAPA Biosystems, Wilmington, MA, #KK2602)], and the thermal cycling was performed as following condition: denaturation at 95°C for 3 min, 5 cycles of denaturation for 20 sec at 98°C, annealing for 15 sec at 65°C and extension for 30 sec at 72°C, followed by a final extension at 72°C for 2 min. Then, 2.5μL of first PCR product was mixed with the 22.5 μL of second PCR mixture [0.35 μM of primers (i5_2nd and i7-BC_mTrbc), and 1x KAPA Hifi Hotstart ReadyMix], and the thermal cycling was performed under the same condition as first PCR. The second-PCR products were purified by an Agencort AM Pure XP kit (Bexkman-Coulter, CA, #A63881) at a 0.7:1 ratio of beads to sample, and eluted with 20 μL of Tris-HCl (pH 8.0). To amplify TCR library and add adaptor sequence for next generation sequencer, the third PCR was performed as follows. 5μL of purified second PCR product was mixed with the 20 μL of third PCR mixture [0.4 μM of primers (i5-BC and i7-BC), and 1x KAPA Hifi Hotstart ReadyMix (KAPA Biosystems, Wilmington, MA, USA, #KK2602)], and the thermal cycling was performed under the same condition as first PCR, excepted for the number of cycles (23 cycles). The third-PCR products were purified as second PCR. The products were pooled and then purified and subjected to dual size selection using ProNex size-selective purification system (Promega) and eluted with 25 μL of Tris-HCl (pH 8.5). Final TCR libraries, whose lengths were about 600 base pairs were sequenced using an Illumina Novaseq 6000 S4 flowcell (67 bp read 1 and 140 bp read 2) (Illumina, USA). Only read2 contained the sequence regarding the definition of T cell clones.

### Data processing of TCR sequencing

Adapter trimming and quality filtering of sequencing data were performed using Cutadapt-3.2 (24) and PRINSEQ-0.20.4 (25). Sequencing data were processed by MiXCR-3.0.5 (26). In MiXCR, Filtered reads were aligned to reference mouse TCR V/D/J sequences registered in the international ImMunoGeneTics information system with the following parameters: -starting-material=rna, −5-end=no-v-primers-, −3-end=c-primers, -adapters=no-adapters, vParameters.geneFeatureToAlign=VTranscript, -vjAlignmentOrder=JThenV. Then, identical sequences were assembled and grouped in clones with PCR and sequencing error correlation with the following parameters: -badQualityThreshold=15, –separateByV=true, –separateByJ=true, -only-productive=true, –region-of-interest=CDR3. The Variable (V) and Joining (J) segment of TCRs were represented in IMGT gene nomenclature.

List of final clones were analyzed by VDJtools-1.2.1 (27). Sequencing reads of sample was normalized to the six times of cell count in each sample by “DownSample” command of VDJtools. T-cell clones were determined as TCR reads with the same TCR V segment, J segment and CDR3 nucleotide sequence. TCR repertoires of divided tumors were pooled other than Figure 3. The processed data have been deposited in NCBI GEO under the accession GSE174225.

### Evaluating the feature of TCR repertoire and the extent of overlap between repertoires

V/J segment usage plots of the TCR repertoire of bilateral tumors were generated by “PlotFancyVJUsage” command of VDJtools. Principle component analysis of V and J segment usage were performed based on the frequency of each V or J segment using prcomp fuction of R (version 3.6.0). The 1 - Pielou index was used to evaluate the clonality of TCR repertoire, which was calculated using the formula: 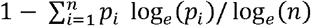 where *p*_i_ is the frequency of clone i for a sample with n unique clones. The Morisita-Horn index was used to estimate the similarity of TCR repertoire between bilateral tumors, which was calculated using the formula:

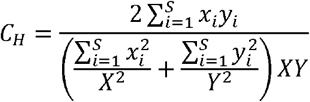

where *x_i_* is the number of clones *i* in the total X reads of one sample, *y_i_* is the number of clones i in the total Y reads of another sample, and S is the number of clones.

The frequency of OL clones between samples is calculated by the geometric mean of the frequencies within each sample.

### Statistical analysis

Statistical analyses were performed using GraphPad Prism (ver8) software (GraphPad Software, La Jolla, CA). Two-sided paired Student’s t-test was run on the comparison of frequency of dLN-Tumor OL clones between ipsilateral and contralateral ones. Ordinary one-way analysis of variance was run to compare the clonality of TCR repertoires between bilateral tumors and between individuals. All other experimental data were analyzed using two-sided unpaired Student’s t-test. Asterisks to indicate significance corresponding to the following: n.s., not significant (P > 0.05), *P ≤ 0.05, **P ≤ 0.01, *** P ≤ 0.001, **** P ≤ 0.0001.

## Acknowledgments

This work was supported by the Japan Society for the Promotion of Science under Grant Number 20281832 and 17929397. We would like to thank Y. Hara for advice in cell sorting, staff of RIBS animal facility for supporting maintenance of animals, member of IGT. Inc., and J. Yasuda for expert technical assistance in TCR repertoire analysis, Editage (www.editage.com) for English language editing.

