## Supplementary material for "T cell receptor (TCR) repertoire analysis reveals a highly conserved TCR repertoire in a bilateral tumor mouse model": Sup Fig1-4, Sup Table 1

**Figure S1**

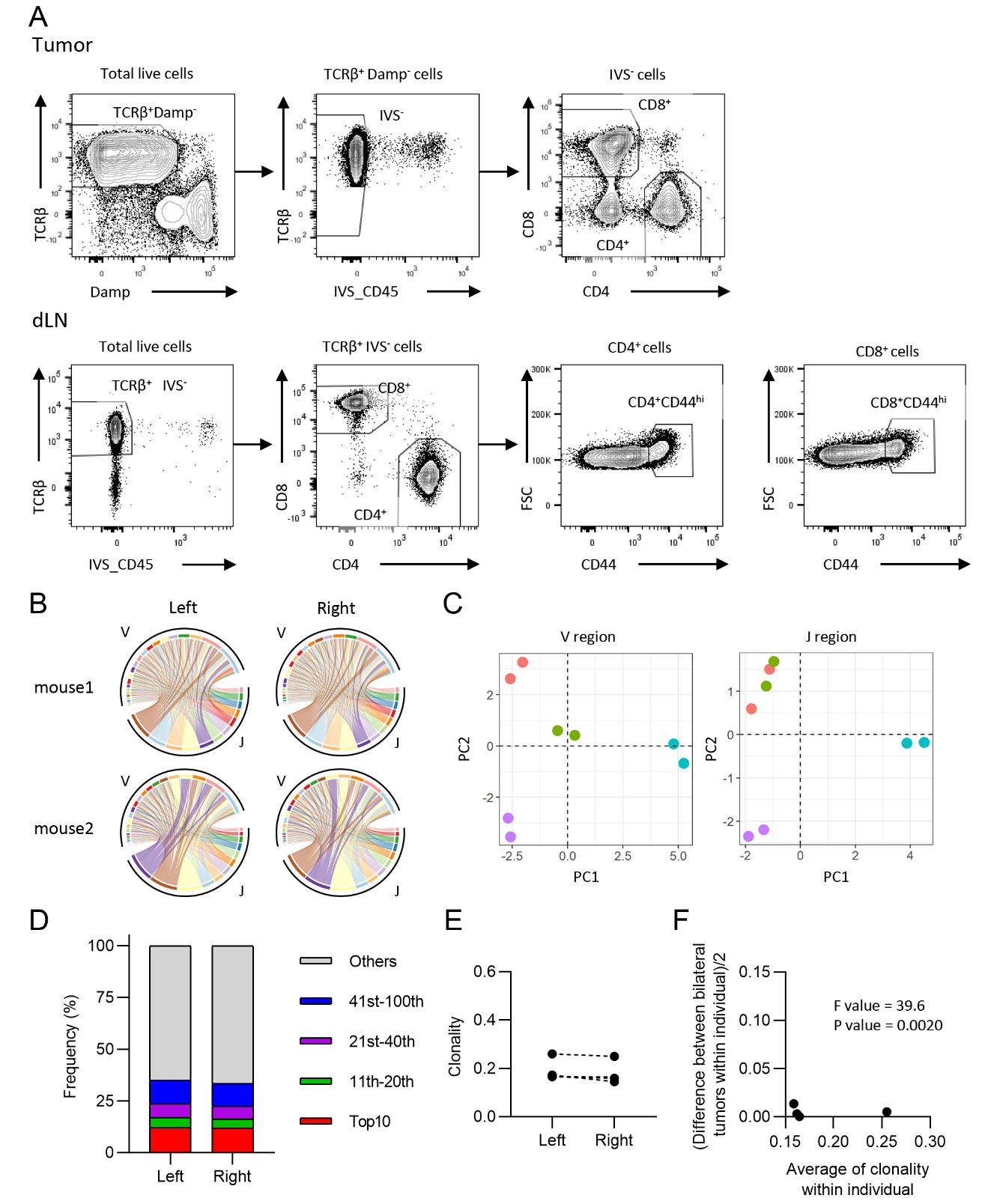

**Fig. S1:** Characteristics of CD4^+^ T cell repertoire in the bilateral tumor. (A) Gating strategy for CD4^+^ and CD8^+^ T cells in the tumor and for CD4^+^CD44^hi^ and CD8^+^CD44^hi^ T cells in the draining lymph node (dLN). (B) V/J segment usage plots of the bilateral CD4^+^ T cell tumor repertoires. Ribbons connecting the V and J segments are scaled by the corresponding V/J pair frequency. (C) Principle component analysis of V and J segment usage of tumor-infiltrating CD4^+^ T cells. Points of the same color indicate left and right tumors of the same individual. (D) Frequency of abundant CD4^+^ T cell clones in the tumor. CD4^+^ T cell clones in the left and right tumor were categorized into five classes based on their rank in each repertoire: Top10, 11th–-20th, 21st–-40th, 41st–-100th, and Others. The total frequency of clones in each class is shown (n = 4). (E) Clonality of the left and right CD4^+^ tumor repertoire. (H) Homoscedasticity plot for variance of clonality in individual mice. The X-axis represents the average clonality of bilateral tumors in each individual. The Y-axis represents variance of clonality between different bilateral tumors within individuals. Sum of squares within mouse = 4.4 × 10^-4^, Sum of squares between mice = 1.3 × 10^-2^, Degree of freedom within mouse = 4, Degree of freedom between mice = 3.

**Figure S2**

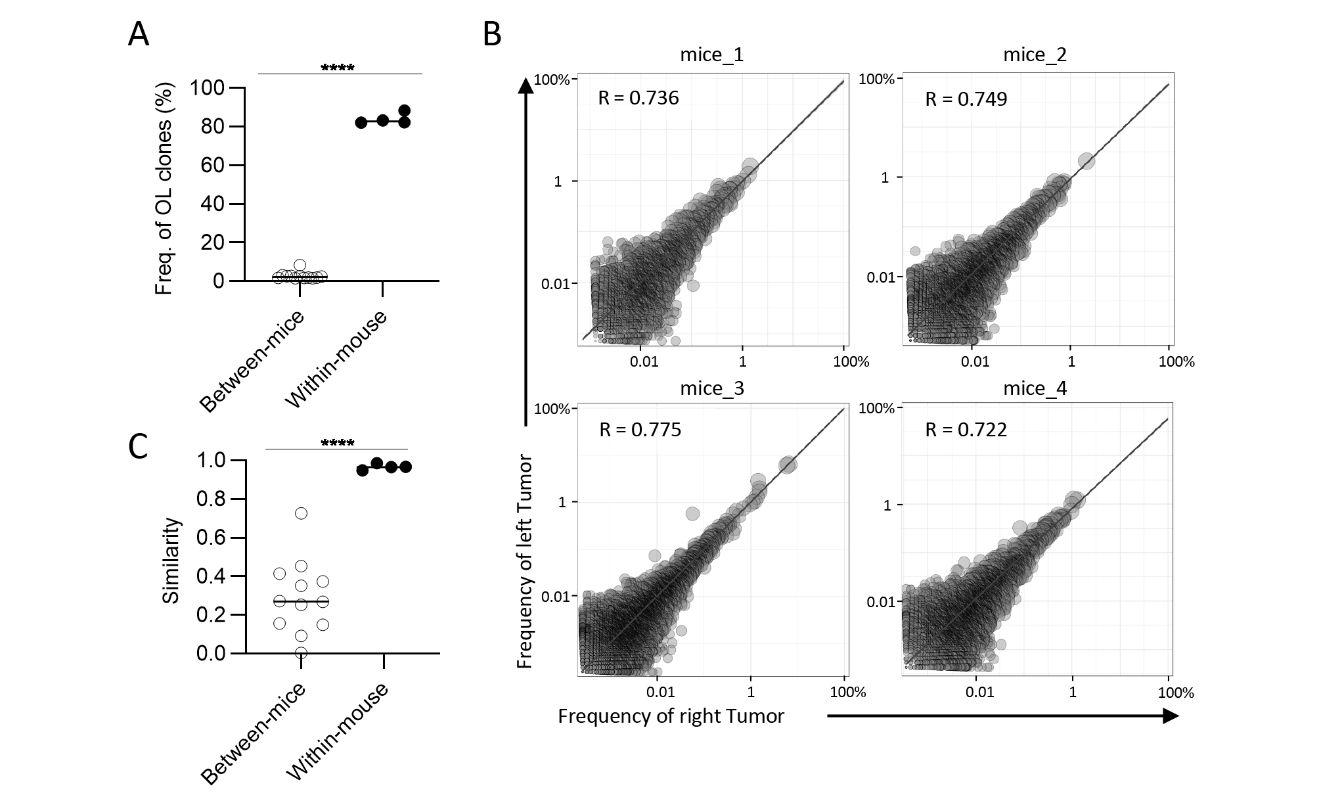

**Fig. S2:** Clonal composition of CD4^+^ T cell repertoires in bilateral tumors. (A) Comparison of the frequency of overlapping clones in tumors between and within mice (between mice, n = 4 × 3; within mice, n = 4). (B) Scatter plot of CD4^+^ T cell clones from bilateral tumors. Each plot represents a single clone with indicated frequency in the left (X-axis) and the right tumors (Y-axis). The dotted line indicates a frequency of 1%. (C) Comparison of the similarity of the tumor repertoires between and within mice. Mean; Two-sided unpaired Student’s t-test (A and C); ****, P ≤ 0.0001.

**Figure S3**

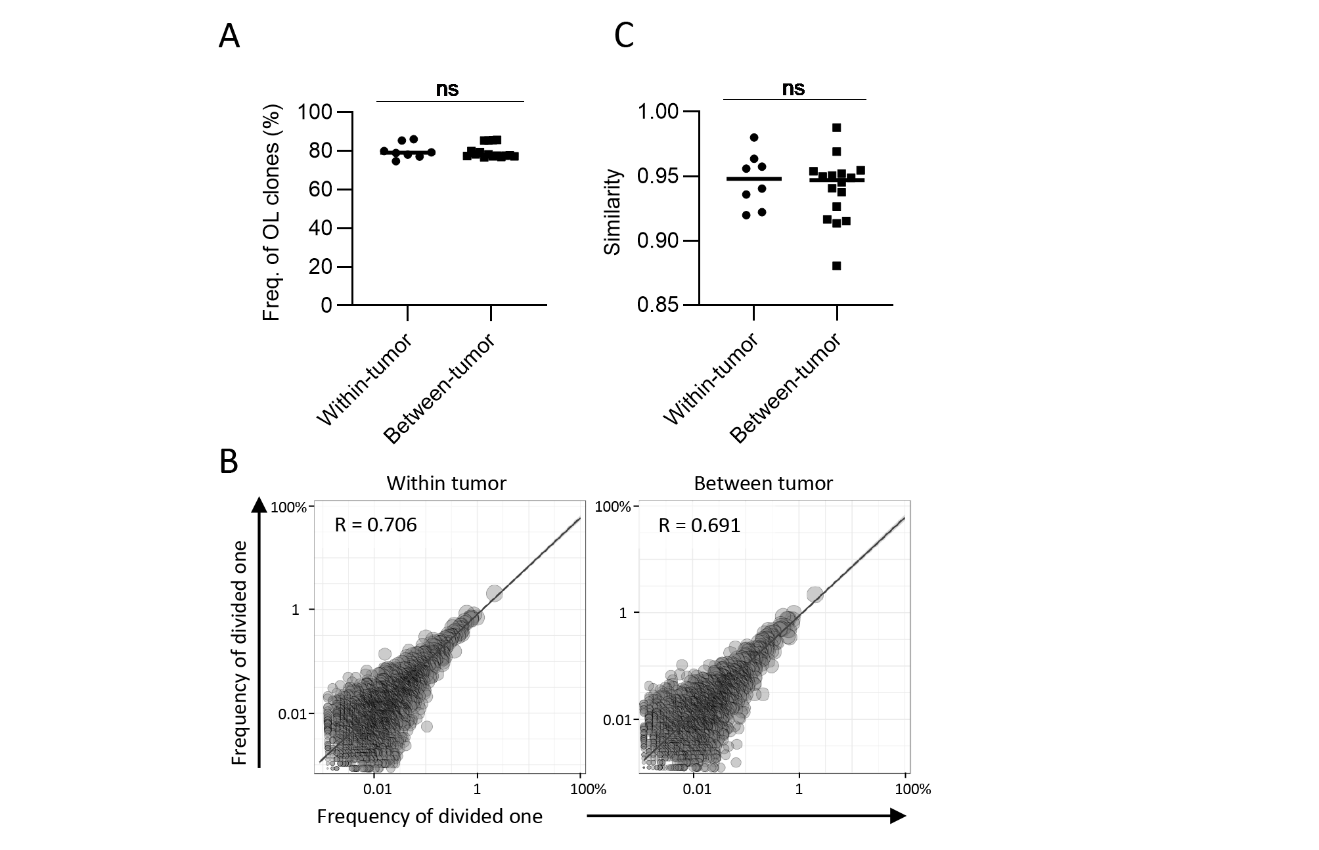

**Fig. S3:** The extent of difference of CD4^+^ T cell repertoires between bilateral tumors. (A) Comparison of the frequency of the clones overlapped between divided same side tumors and bilateral tumor fragments (n = 4). (B) Scatter plot of CD8^+^ T cell clones from bilateral tumors. Each plot represents a single clone with indicated frequency in each tumor fragment. The dotted line indicates a frequency of 1%. (C) Comparison of the similarity of the tumor repertoires between divided same tumors and bilateral tumor fragments. (A and C) Within-tumor, n = 4 × 2; between-tumor, n = 4× 4; Mean; Two-sided unpaired Student’s t-test; ns, non-significant.

**Figure S4**

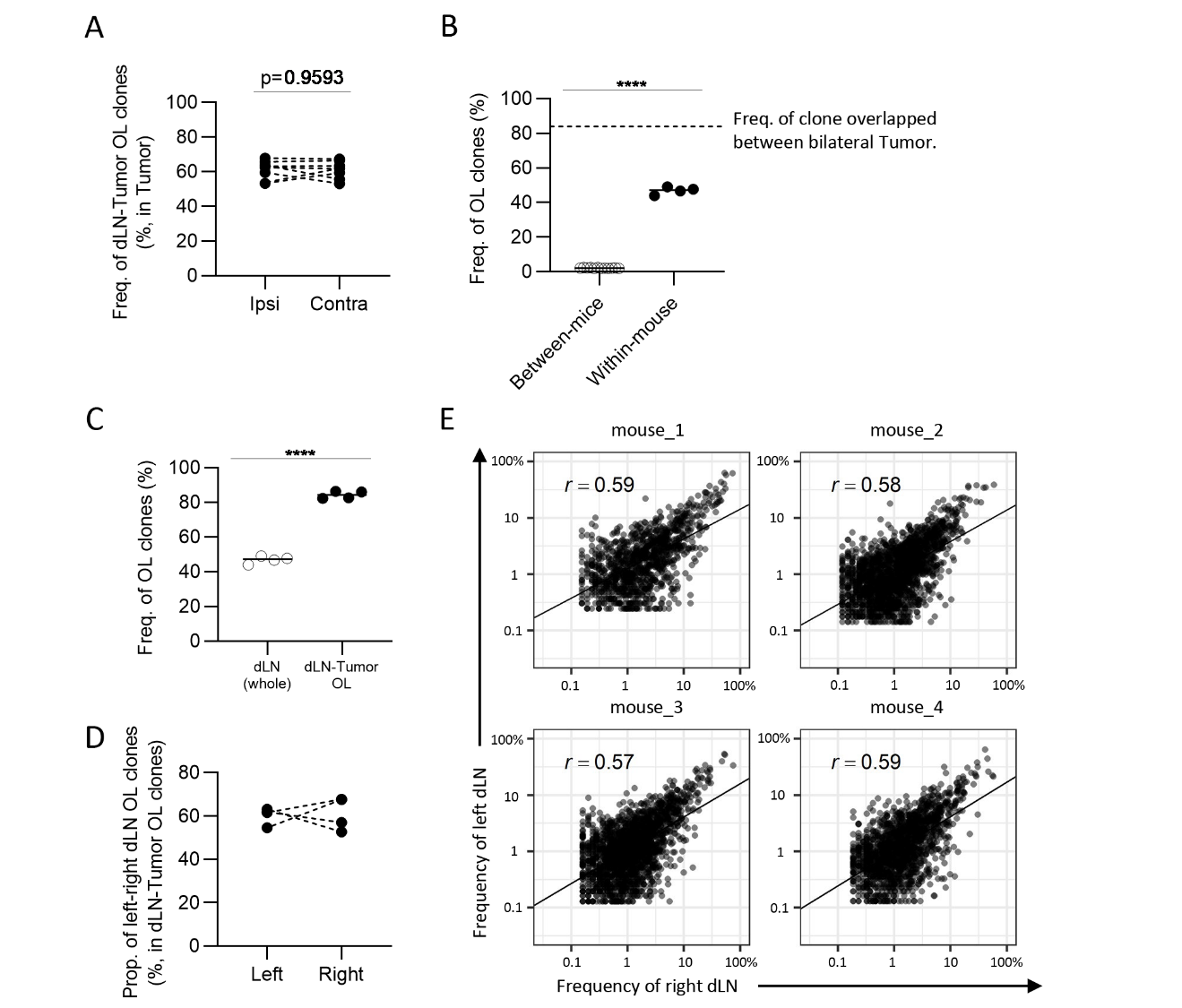

**Fig. S4:** The similarity of CD4^+^ T cell repertoires in the bilateral dLNs. (A) The frequency of the clones overlapped between the dLN and its ipsilateral and contralateral tumor. Ipsi, ipsilateral; Contra, contralateral (n = 4 × 2). (B) Comparison of the frequency of overlapping clones in dLN between and within mice (between mice, n = 4 × 3; within mice, n = 4). Dotted line indicates the frequency of clones overlapped between bilateral tumors within mice. (C) Comparison of the frequency of left-right overlapping clones within whole dLNs and dLN-tumor overlapping repertoire (n = 4). (D) The proportion of left–-right overlapping clones within the dLN-tumor overlapping repertoires. (E) Scatter plot of left–-right overlapping clones within the dLN-tumor overlapping repertoires. Each plot represents a single clone with indicated frequency in each dLNs. Mean; Two-sided unpaired Student’s t-test; ns, non-significant. **** P ≤ 0.0001.

**Table S1**

|  | antigen | conjugate | company | clone |
| --- | --- | --- | --- | --- |
| Tumor | CD4 | BV510 | BD | RM4-4 |
|  | CD8a | APC | BioLegend | 53-6.7 |
|  | PD-1(CD279) | PE-Cy7 | BioLegend | 29F.1A12 |
|  | TCRβchain | APC-cy7 | BioLegend | H57-597 |
|  | Ly108 | BB700 | BD | 13G3 |
|  | Tim3 | BV421 | BioLegend | RMT3-23 |
|  | CD11b | PE | BioLegend | M1/70 |
|  | CD45R(B220) | PE | BioLegend | RA3-6B2 |
|  | NK1.1 | PE | eBioscience | PK136 |
|  | TER119 | PE | BioLegend | TER-119 |
| dLN | CD4 | Pacific Blue | BioLegend | RM4-4 |
|  | CD8a | APC | BioLegend | 53-6.7 |
|  | CD44 | PerCP_Cy5.5 | BioLegend | IM7 |
|  | TCRβchain | APCCy7 | BioLegend | H57-597 |
|  | CD11b | Biotin | BioLegend | M1/70 |
|  | CD45R(B220) | Biotin | BioLegend | RA3-6B2 |
|  | NK1.1 | Biotin | BioLegend | PK136 |
|  | Ly6G | Biotin | BioLegend | 1A8 |

Table S1: Summary of antibodies used in this study, listing the target protein, conjugated fluorophore, manufacture, antibody clone.
